# Flexible multiplane structured illumination microscope with a four-camera detector

**DOI:** 10.1101/2020.12.03.410886

**Authors:** Karl A. Johnson, Daniel Noble, Rosa Machado, Guy M. Hagen

**Affiliations:** UCCS BioFrontiers Center, University of Colorado Colorado Springs, 1420 Austin Bluffs Parkway, Colorado Springs, Colorado 80919, USA

## Abstract

Fluorescence microscopy provides an unparalleled tool for imaging biological samples. However, producing high-quality volumetric images quickly and without excessive complexity remains a challenge. Here, we demonstrate a simple multi-camera structured illumination microscope (SIM) capable of simultaneously imaging multiple focal planes, allowing for the capture of 3D fluorescent images without any axial movement of the sample. This simple setup allows for the rapid acquisition of many different 3D imaging modes, including 3D time lapses, high-axial-resolution 3D images, and large 3D mosaics.

## 1. Introduction

Structured illumination microscopy (SIM) is a technique in fluorescence microscopy in which sets of images are acquired with shifting illumination patterns; subsequent image processing of these image sets can yield results with optical sectioning, resolution beyond the diffraction limit (super-resolution), or both [1–6]. Since its emergence over two decades ago [7], SIM has matured significantly as an imaging technique, with multiple proposed methods for generation of the SIM pattern [4,6–11] as well as for processing of the image data [3,6,12–14]. Compared to other super-resolution techniques, the speed, high signal-to-noise ratio, and low excitation light intensities characteristic to SIM make it ideal for the imaging of live samples in 3D or 4D.

Producing a 3D image with fluorescence microscopy requires intensity information throughout the sample to later be localized to a specific location in 3D space. With SIM, this is traditionally accomplished by acquiring multiple optically sectioned images of the sample while translating the sample axially though the focal plane of the microscope. This simple method immediately yields discrete XY slices of the sample, which can then be stacked into a 3D image. Though this sequential approach remains a staple of fluorescence microscopy, the substantial light exposure to the sample, long acquisition times, and possible agitation to the sample incurred by this method are significant drawbacks.

As such, several other methods have been developed as an alternative to this traditional approach. Prominent among these is multifocal plane microscopy, in which multiple focus-shifted image planes are imaged simultaneously, removing the need for sample movement. This can be achieved using a variety of techniques, and in recent years multiple studies have demonstrated several methods to image multiple image planes side-by-side onto a single detector. The two most popular approaches to achieve this involve either the use of a multi-focal grating (MFG) [15–19] or a variable path length image splitting prism [20,21]. Though these techniques can produce high-quality results, the highly specialized optical elements central to these approaches are difficult to design and can restrict the versatility of the resulting setups. Since both approaches have a defocus distance defined by the physical features of the specialty optics being used, the distance between imaging planes in the sample cannot be easily adjusted. Additionally, both of these optical systems are designed to be used with a particular objective, and as such switching between objectives may compromise image quality. These substantial limitations in the ability to change slice spacing and objectives limit the adaptability of such systems for everyday use.

Recently, Xiao et. al. proposed a reconfigurable multiplane microscope which uses an array of beam splitters to image multiple focal planes on a single detector [23]. Unlike the aforementioned methods, this approach is compatible with a variety of microscope objectives, due to its comparative optical simplicity. Still, this approach requires the construction of a moderately complex ‘z-splitter’ assembly, and the separation between image planes produced by this optical system cannot be modified without changing critical dimensions of the ‘z-splitter.’

Multi-camera approaches have also previously been explored in fluorescence microscopy as a method for multifocal plane microscopy. These methods are particularly attractive due to their optical simplicity and versatility. In 2004, Prabhat et. al. proposed an optical system in which two cameras are placed at different focal positions relative to the microscope’s tube lens, resulting in each camera capturing an image at a different focal plane in the sample [24]. Others have since expanded upon this idea by using this multidetector method for 3D localization microscopy [25] or by expanding the number of detectors to three or four [25,26].

Unfortunately, the viability of multi-camera imaging has been limited by the high cost of scientific cameras for many years. However, recently, complementary metal oxide semiconductor (CMOS) sensor technology has improved significantly in both performance and price, yielding a variety of affordable cameras with sufficient sensitivity and noise performance for fluorescence microscopy. In 2018, Babcock demonstrated a 3D localization microscopy system using four industrial-grade CMOS cameras [25]. The cameras could detect the extremely dim signals necessary for localization microscopy (10-1000 photons/pixel) while costing an order of magnitude less than typical scientific CMOS (sCMOS) cameras.

Here, we demonstrate three different imaging modes achievable using a variation of Babcock’s detection system in conjunction with super-resolution SIM methods. Switching between modes can be performed without physical manipulation of the optical setup. This allows for versatility in imaging, as different types of samples are best imaged by different modes. As live cells constantly move, they are best imaged simply by taking a sequence of 3D images over time, yielding a super-resolution, 3D movie of the cell. For static samples with significant axial extent, a Z-stack of 3D images can be acquired, producing a high-axial-resolution and large 3D image but with a four-fold reduction in sample movements. Finally, larger samples can be imaged by taking a mosaic of 3D images, resulting in images with a large field of view (FOV) and useful axial information. Image processing for all these imaging modes can be performed using our open source software for SIM (SIMToolbox) [27] along with standard tools in MATLAB (The MathWorks, Natick, MA) and ImageJ [28].

## 2. Materials and Methods

### 2.1 Microscope set-up and data acquisition

This project uses a home-built SIM set-up based on the same design as described in our previous publications [8,12,29,30]. The SIM system is based on an IX83 microscope (Olympus, Tokyo, Japan) with our 4-camera setup serving as a detector. The data in this study were collected using the UPLSAPO 30×/1.05 NA silicone oil immersion and UPLSAPO 60×/1.35 NA oil immersion objectives (Olympus), though our setup is compatible with any objective meant to be used with the IX83 microscope. Sample movement was controlled with an ASI motorized XY, piezo Z stage (Applied Scientific Instrumentation, Eugene, Oregon). We used a quadruple band fluorescence filter set (part 89000, Chroma, Bellows Falls, VT). To synchronize the four cameras with SIM illumination and stage movement, we used Andor IQ software (Belfast, Northern Ireland, UK).

Our SIM system uses a ferroelectric liquid crystal on silicon (LCOS) microdisplay (type SXGA-3DM, Fourth Dimension Displays, Dalgety Bay, Fife, UK). The LCOS microdisplay has been utilized before in SIM and related methods in fluorescence microscopy [2,8,12,27,31–34] and allows patterns of illumination to be projected on to the sample that can be reconfigured by changing the image displayed on the device. The light source (Lumencor Spectra-X, Beaverton, OR, USA) is toggled off between SIM patterns and during camera readout.

The four cameras used in our set up are the Blackfly-S USB 3 (model: BFS-U3-31S4M, FLIR, Arlington, VA). The manufacturer’s specifications for relevant parameters of this camera are shown in Table 1. These cameras have desirable parameters for fluorescence microscopy such as low noise, but at a much lower price point than the sCMOS cameras typically used.

**Table 1.** Camera parameters

|  |  |
| --- | --- |
| **Camera** | FLIR Blackfly S |
| **Part number** | BFS-U3-3154M |
| **Sensor type** | Sony IMX265 |
| **Quantum efficiency** | 62% at 525 nm |
| **Read noise** | 2.26 $e^-$ |
| **Pixel size** | 3.45 $\mu\text{m}$ |
| **Format** | 2048×1536 |
| **Maximum frame rate** | 55 FPS |
| **Readout method** | global shutter |
| **Sensor size** | 7.18×5.32 mm (1/1.8" format) |

The four cameras are mounted on XYZ translation stages (MT3, ThorLabs) to enable precise positioning of each camera. SpinView software (FLIR) was used to interface with the cameras and to acquire the images. Fig. 1 shows a simplified diagram and a to-scale model of the optical system.

**Fig. 1.**
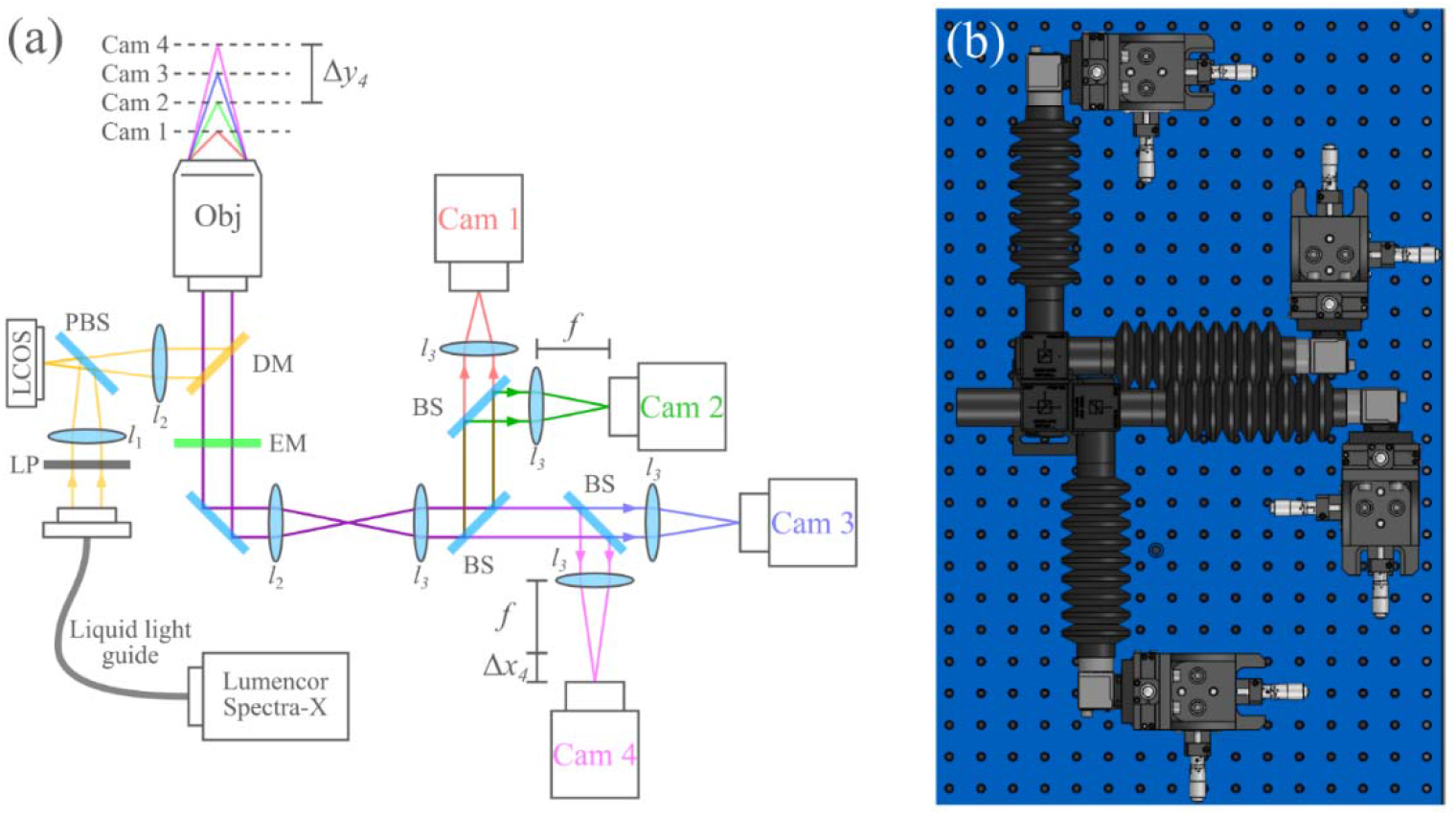
(a) Overview of the optical system. Δx_4_ and Δy_4_ indicate the sensor defocus and sample defocus of camera 4, respectively. LP, linear polarizer; LCOS, liquid crystal on silicon microdisplay; PBS, polarizing beam splitter; DM, dichroic mirror; EM, emission filter; BS, 50-50 beam splitter; l_1_, 50mm FL; l_2_, 180mm FL Olympus tube lens (part SW-TLU); l_3_, 175mm FL. (b), to-scale schematic of the 4-camera detection system.

In order to image multiple distinct planes in the sample, each camera was placed at a slightly different distance from each camera’s respective relay lens (l_3_ in Fig. 1). Importantly, the light paths of all four cameras are based on a 4-f design. This design ensures near telecentricity on all four cameras, and as such, defocusing the detectors results in nearly no change in magnification. To position the cameras appropriately, we first aligned all cameras at the focal length of each camera’s respective l_3_. Then, each camera was displaced using its translation stage to achieve a particular defocus. The relationship between this image sensor defocus Δ*x*_*n*_ and the defocus of the sensor conjugate (focal plane) in the sample Δ*y*_*n*_ for each camera *n* can be described by Eq. 1., where *M* is the magnification of the optical system and η_sample_ is the refractive index of the sample.

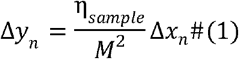

In our setup, we positioned each camera such that the focal planes were evenly spaced by a slice spacing Δ*s*. Additionally, to reduce aberrations caused by excessive defocus distances from the focal plane of the SIM pattern, cameras were defocused both forwards and backwards, with camera 2 kept stationary. This yields the following formulae (Eqs. 2-5) for the translation stage displacements of each camera, given the desired slice spacing Δ*s*:

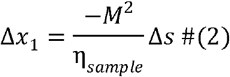

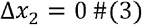

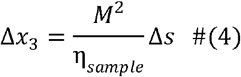

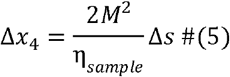

To verify Eq. (1), a 45nm fluorescent bead (f10720, Molecular Probes, Eugene, OR) dried on a coverslip and overlaid with microscope oil (η = 1.518) was imaged under widefield illumination with all four cameras and a 60×/1.35 NA oil immersion objective. To acquire volumetric data of the bead images on each camera, the z stage was scanned over 3um in 20nm increments. The orthogonal projections of the resulting data, shown in Fig. 2, indicate that the bead’s images are displaced axially by an amount close to that predicted by (1). Here, each camera was displaced in 1.5 mm increments, and the effective magnification of the system was (180mm/175mm) · 60× = 61.7× due to the re-imaging system. Using these figures with equation (1) yields a predicted slice increment of 630nm in the sample. The slight discrepancy between the measured displacement and the predicted displacement can be attributed to misalignments in the optical system, which effect the telecentricity and therefore the effective magnification of the system for each camera.

**Fig. 2.**
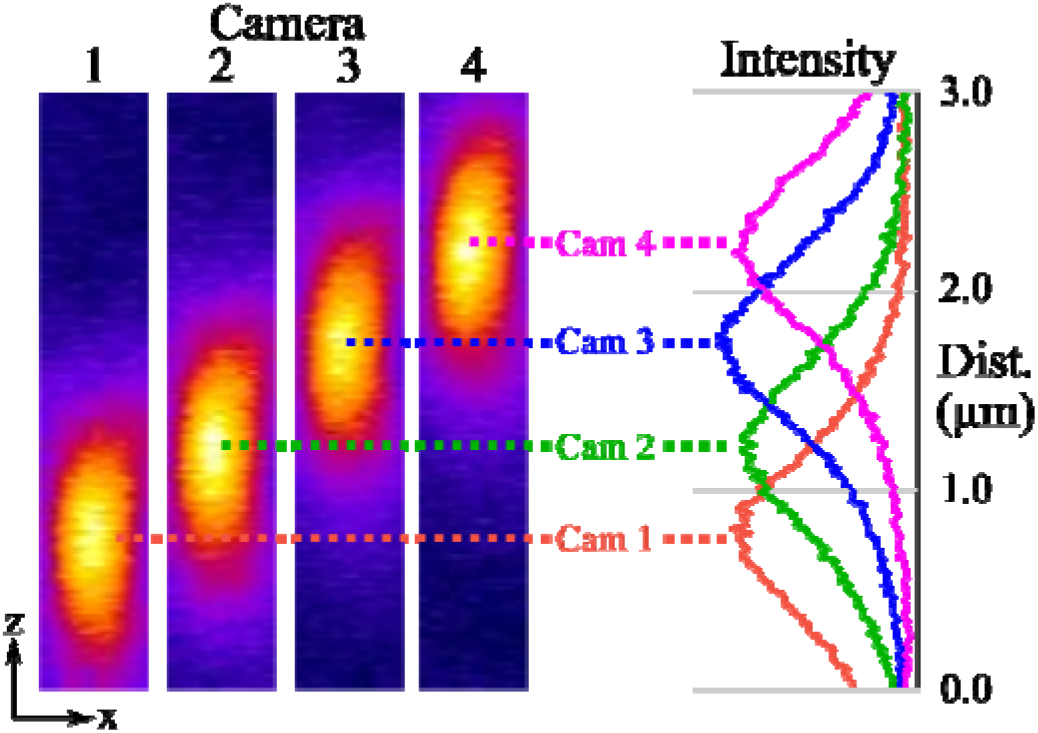
45nm fluorescent nanobead imaged with defocused cameras and widefield illumination. The images to the left of the figure show the orthogonal projection of the image stacks captured by each camera. The plot to the right, which has the same z scale as the images, shows overlaid intensity plots of each image along the z axis. In this data, the cameras were defocused with a step of 1.5mm at a magnification of 60x. The measured average displacement between the cameras’ focal planes aligns somewhat well with the theoretical displacement of 630nm.

### 2.2 Cell lines and reagents

Hep-G2 cells were maintained at 37°C and 100% humidity in DMEM supplemented with 10% fetal bovine serum, 100 U/mL penicillin, 100 U/mL streptomycin, and L-glutamate (Invitrogen).

### 2.3 Preparation of samples for imaging

Hep-G2 cells were grown in coverslip-bottom imaging dishes for 48 hours, then labeled with Mitotracker Red CMXROS (M7512, Molecular Probes) according to the manufactures recommended protocol. Briefly, cells were labeled with 1mM Mitotracker for 30 minutes at room temperature, then washed twice with and imaged in phosphate-buffered saline, pH 7.4.

We also imaged 3^rd^ instar *Drosophila melanogaster* (ppk-CD4-tdTom) larvae, which express CD4-tdTomato in the sensory neurons. We anesthetized the larvae using isoflurane. The larvae were then placed on a chamber slide with a drop of SlowFade Gold antifade mountant (Thermo Fisher, S36936), then sealed with a #1.5 coverslip.

Finally, we imaged a GFP labelled mouse brain sample acquired from SunJin Lab (Hsinchu City, Taiwan). This sample is a 250um thick coronal section which was cleared and mounted by SunJin Lab using the RapiClear 1.52 reagent.

### 2.4 Image pre-processing

Immediately after acquisition, hot pixels appear throughout most of the raw image data, as the industrial CMOS cameras used in this study are not actively cooled and were operated at room temperature. Since hot-pixels can cause undesirable artifacts in our super-resolution SIM reconstruction algorithm, hot-pixel removal is necessary prior to SIM reconstruction. To do this, we collected dark images with each of the cameras using the same exposure settings as in the image data, then subtracted these dark images from each frame of the raw data. While this simple method is only effective on the hot pixels which appear in both the dark frames and the raw data, these hot pixels make up the vast majority of the total number of hot pixels in the raw data, and as such this method is satisfactory.

### 2.5 SIM data processing

SIM reconstructions were performed as previously described using SIMToolbox, an open source and freely available program that our group developed for processing SIM data [27]. We generated optically sectioned, enhanced resolution images using an established Bayesian estimation method, maximum *a posteriori* probability SIM (MAP-SIM) [12,29,30]. Wide-field (WF) and conventional resolution, optically sectioned (OS-SIM) images were also reconstructed using SIMToolbox.

### 2.6 Image Registration and Assembly/Stitching

After SIM reconstruction, it is necessary to digitally align the images from the four cameras, as it is infeasible to perfectly align the images from each camera using the translation stages. First, a single processed frame is selected from each camera to perform image registration. After selecting one of the cameras as a reference, all other images are registered to the reference camera’s image using a gradient descent optimization method. This registration process yields an alignment matrix for each camera which describes the transformation necessary to transform each camera’s image to align with the reference camera’s image. Note that in case of significant defocus between cameras, single frames may vary too greatly between cameras for registration to be performed. In this case, the registration process can be performed on a maximum intensity projection of each camera’s data, or reference images of a calibration sample, instead of a single processed frame. Next, all processed images from the non-reference cameras are simply transformed using the alignment matrices found during registration, yielding images that are aligned to the reference. This image registration and transformation process was performed using the image processing toolbox for MATLAB.

After alignment of the images from each camera, the data is organized into four separate image stacks (one for each camera). The process to reconfigure this data into its final form is dependent on the imaging mode. For 3D time-lapse imaging, the four image stacks can simply be regrouped into several image stacks, one for each time position. For Z stacks of 3D images, interlacing the four image stacks into a single image stack produces the final 3D image. To generate 3D mosaics, the four camera image stacks must be first regrouped into several image stacks, each containing one 3D image tile. Similar to the time-lapse mode, each of these image stacks will contain four slices, one from each camera. These 3D images can then be stitched into the final image. We performed image stitching using Preibisch’s plugin for ImageJ [35]. This entire data processing procedure is summarized in Fig. 3.

**Fig. 3.**
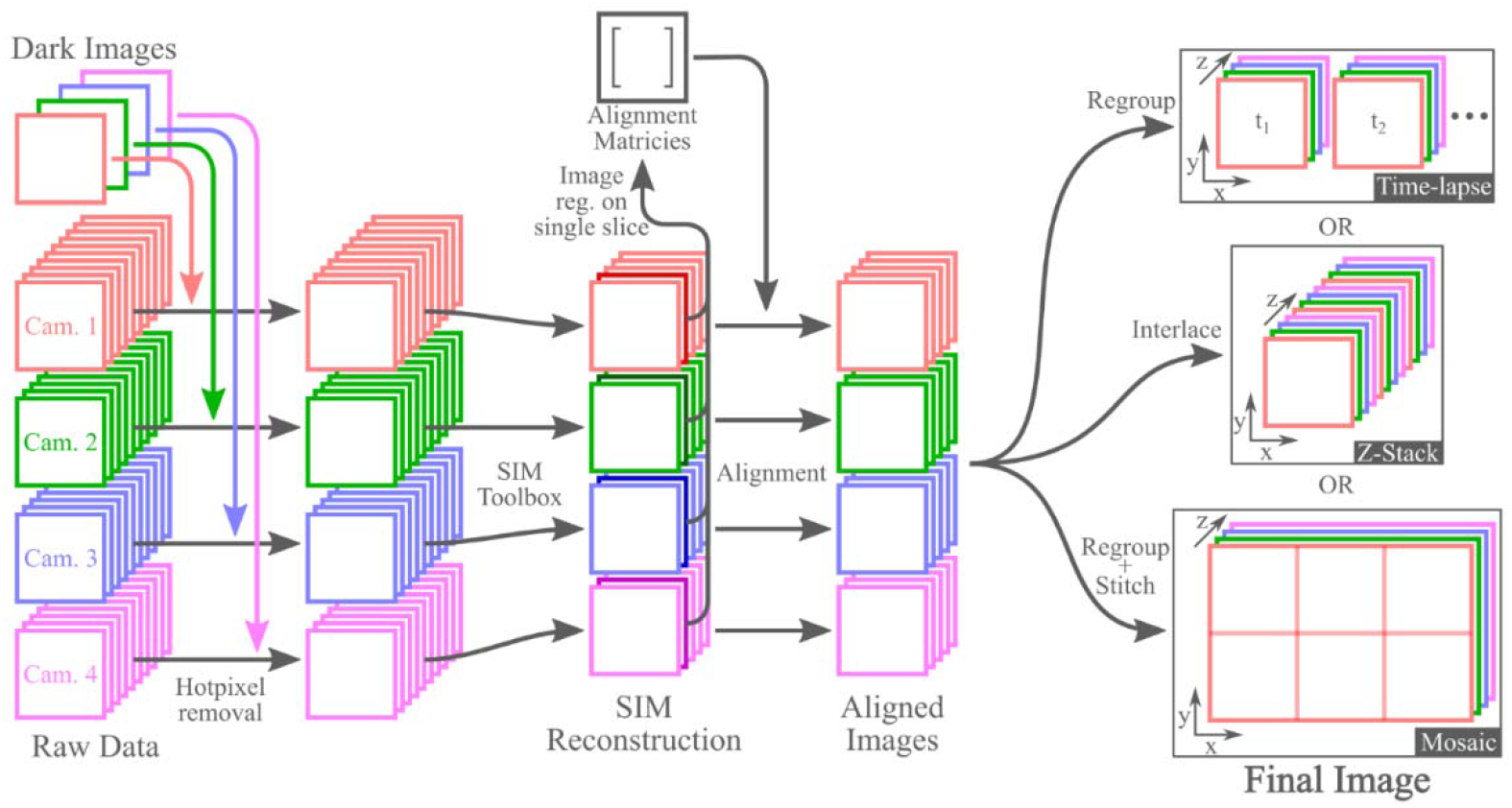
Overview of the standard processing procedure for all three imaging modes. Note that the hot pixel removal and image registration steps only must be performed once per data set, and do not have to be repeated when doing processing for each subsequent SIM reconstruction method. Additionally, while this figure shows the image registration as being performed on a single slice of the reconstructed SIM data, these are not the only images that registration can be performed on. Registration can also be performed on a maximum intensity projection of each camera’s reconstructed data, or on images of a calibration slide, as discussed in the text.

## 3. Results

To demonstrate the time-lapse imaging mode of our system, we imaged mitochondrial dynamics in HEP-G2 cells over three minutes. 3D images were acquired at three second intervals, resulting in 60 total frames, four of which are shown in Fig. 4 (see Visualization 1 for a video containing all frames of the time-lapse). As evident in the figure, MAPSIM improves the resolution compared to the WF reconstruction and eliminates out of focus light. Due to the use of the four-camera detection system, all images were acquired with no physical z-movement of the sample – and, with our SIM system, no moving parts at all during acquisition. To display the 3D information in Figs. 4, 5 and 6, the images were depth coded using the isolum lookup table [36].

**Fig. 4.**
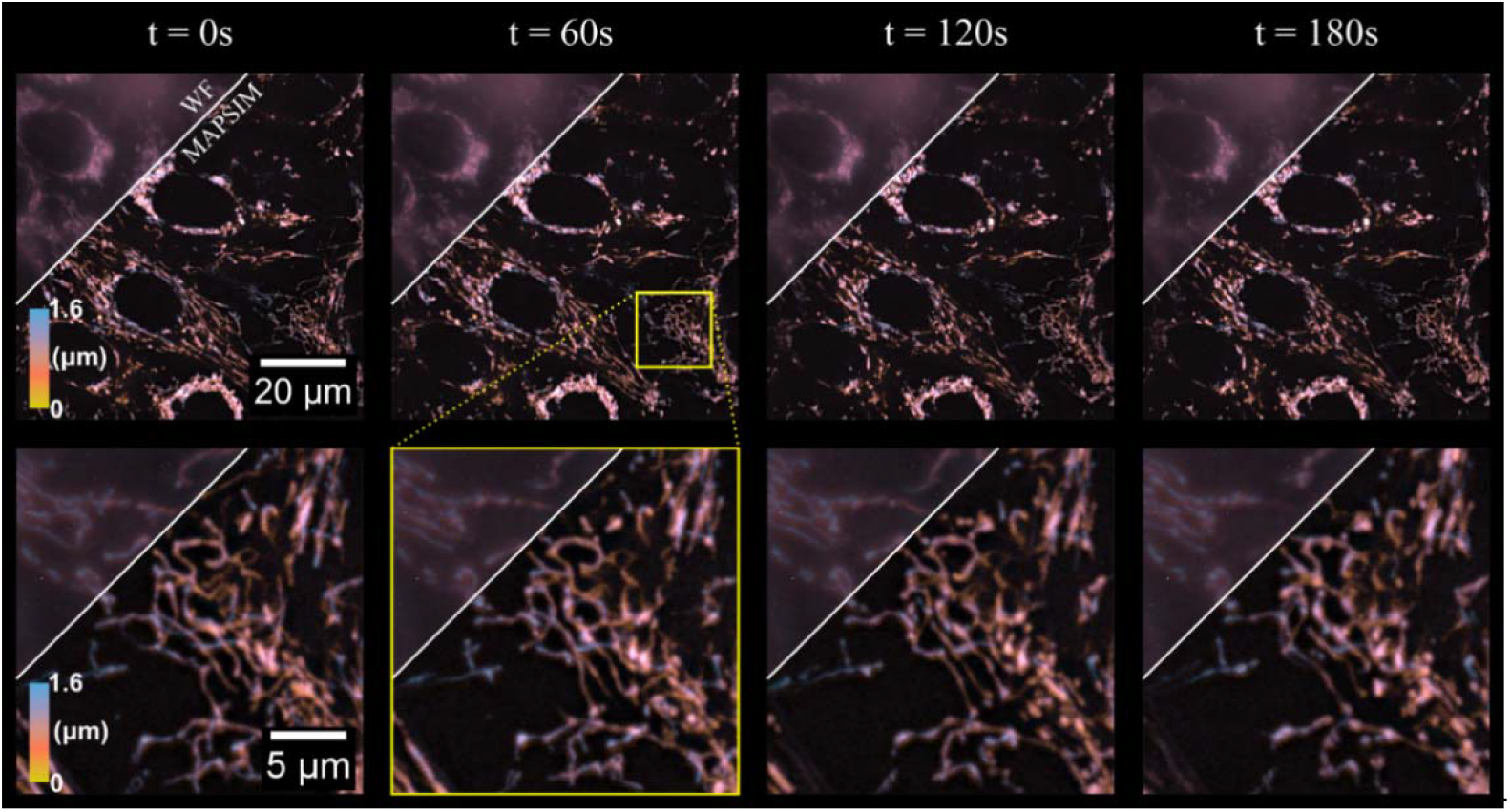
3D time-lapse of mitochondrial dynamics in HEP-G2 cells. The region above the diagonal line on each frame shows the WF reconstruction of each time frame, with the region below the line showing the MAPSIM reconstruction of the data. Objective: UPLSAPO 60X/1.35 NA oil immersion. Imaged at 37 °C and 5% CO2 using type 37 oil (Cargille).

**Fig. 5.**
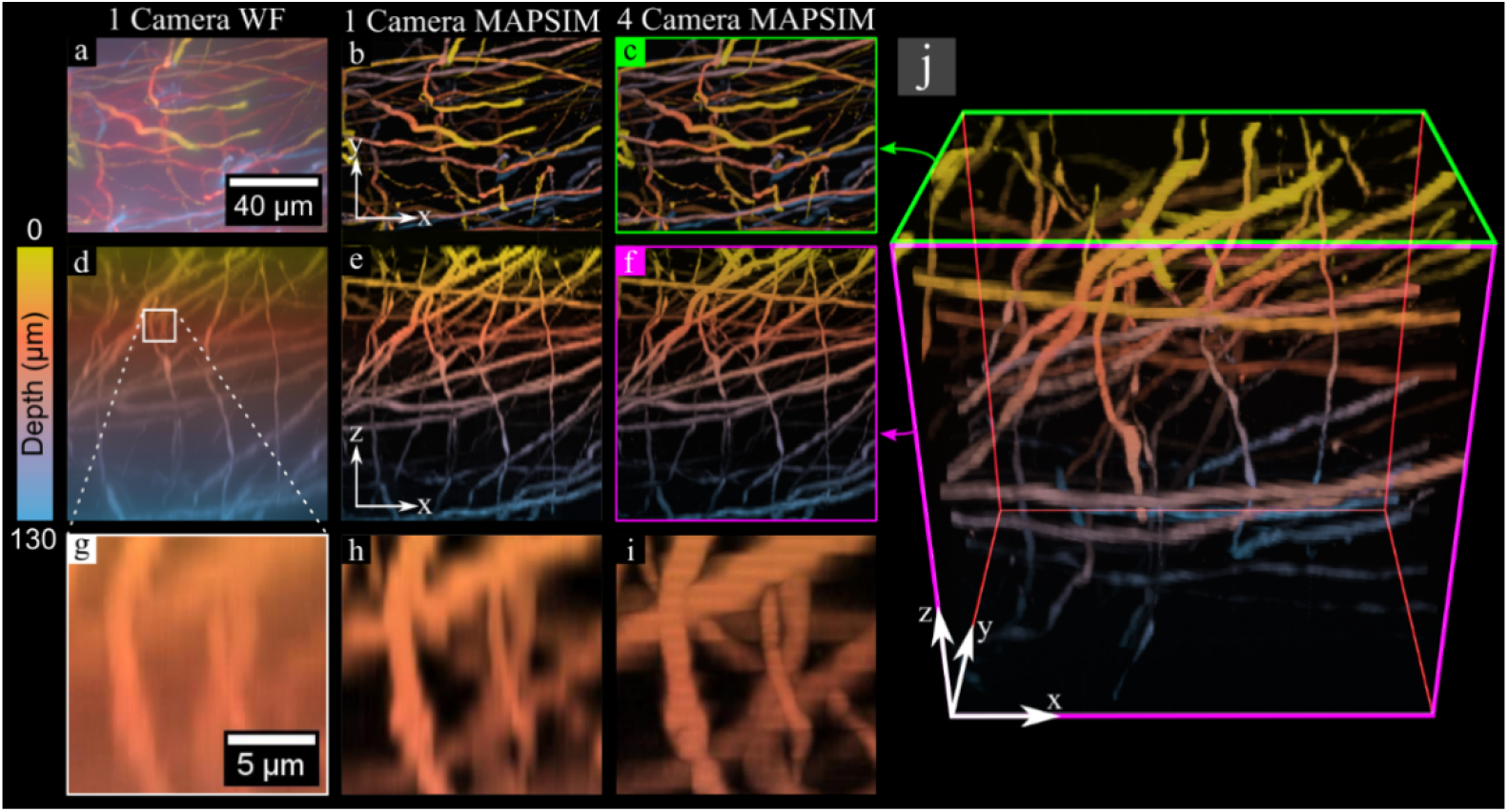
3D image of neurons in a GFP-labelled mouse brain, acquired using the Z-stack imaging mode. All data is color-coded according to the scale bar at the left of the figure. (a), (b), and (c) show X-Y maximum intensity projections of the sample, with (a) showing the WF reconstruction from a single camera, (b) the MAPSIM reconstruction from one camera, and (c) the MAPSIM reconstruction from all four cameras. (d), (e), and (f) show X-Z projections of the same data, respectively. (g), (h), and (i) show a zoom-ins of the X-Z projection from the region highlighted in (d). (j) shows a 3D rendering of the 4-camera MAPSIM reconstruction of the dataset, with the perspectives shown in (c) and (f) highlighted in green and magenta, respectively. Objective: UPLSAPO 60X/1.35 NA oil immersion.

**Fig. 6.**
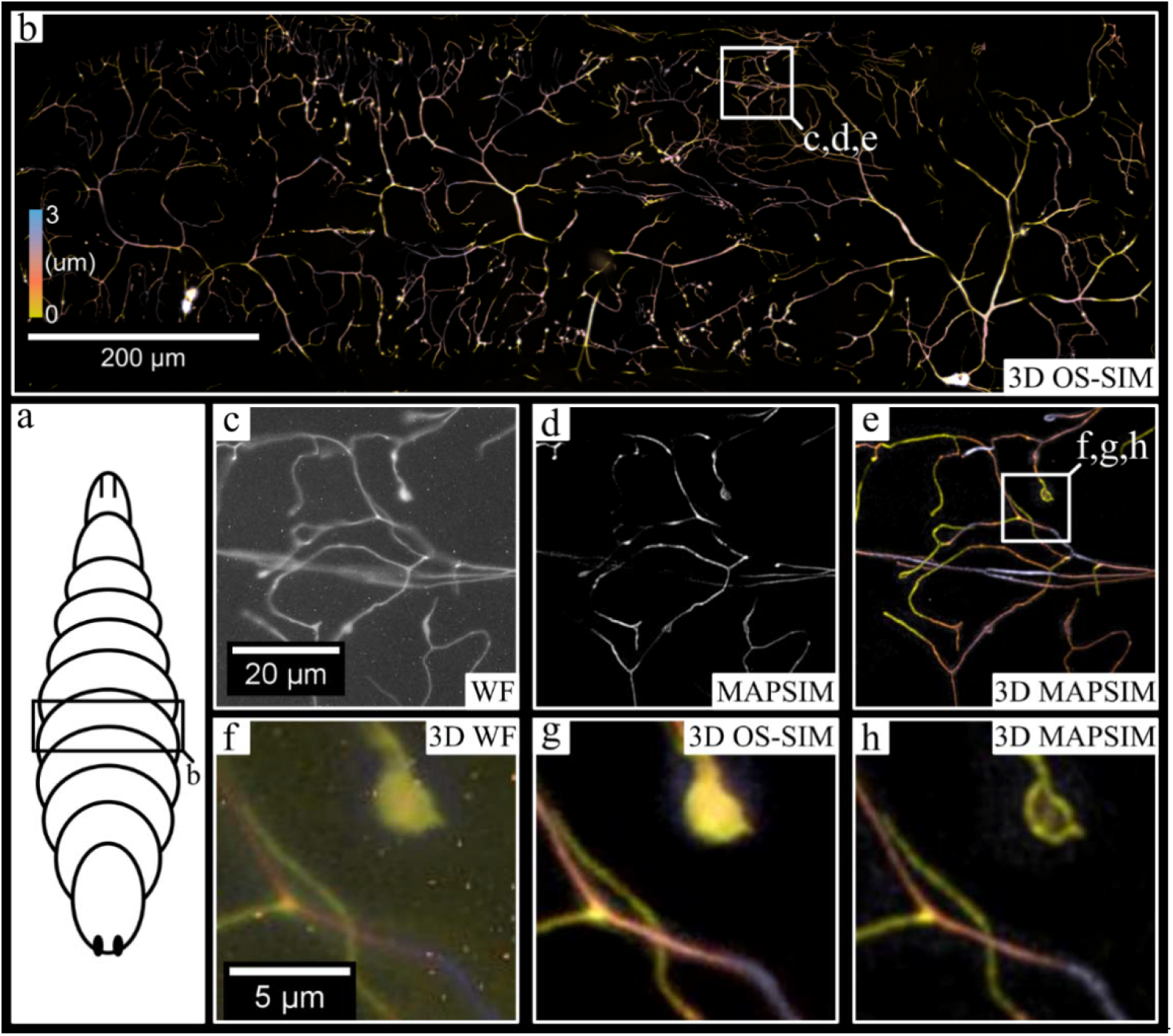
Large FOV, 3D imaging of neurons in an abdominal segment of a Drosophila melanogaster larvae. (a) Generic illustration of a drosophila larvae, with the approximate imaging region labeled. (b) shows the entire 3D, stitched image produced from a mosaic of 3D images generated using the multi-camera system. (c), (d), and (e) show the region indicated in (b), with (c) showing the WF reconstruction from a single camera, (d) showing the MAP-SIM reconstruction from a single camera, and (e) showing the color-coded 3D image produced from MAP-SIM reconstruction using all four cameras. (f), (g), and (h) show a comparison between SIM reconstruction methods on the region indicated in (e), using all four cameras: (f), WF; (g), OS-SIM; (h), MAP-SIM. Note that hot-pixel removal was not performed on the images in (c) and (f) for demonstrative purposes. Objective: UPLSAPO 30x/1.05 NA silicone oil immersion.

Next, we acquired a 130-um thick 3D image of neurons in a GFP-labelled mouse brain sample at 60× magnification to demonstrate the 3D Z-stack imaging mode. Though this 3D image has a size of 2048×1536×316, it was acquired in under two minutes. Due to the ability to capture four z-planes at once with the four-camera setup, the 316 z-planes in this image were acquired with just 79 movements of the z-stage. This image has ∼416 nm sampling in Z and 57.5nm sampling in XY.

Finally, we demonstrated the 3D mosaic imaging mode by imaging CD8-tdTomato labelled neurons in *Drosophila melanogaster* larvae (Fig. 6). The data which produced the final 6957×10151-pixel (∼800×1150×3 um) image was acquired in under 10 minutes. For visualization purposes, this image was rotated, cropped, and color-coded to the region (∼1000×300×3 um) shown in Fig. 6b. As shown in Fig. 6d, the MAPSIM data from a single camera alone shows improved resolution over the widefield reconstruction (Fig. 6c), but MAPSIM’s optical sectioning prevents accurate visualization of neurons that deviate from the image’s narrow depth of field. When all four cameras are included (Fig. 6e), the data not only provides useful depth information, but also has a larger effective depth of field than that of a single z-plane. This ability for the four-camera system to enhance the axial FOV of the montage substantially reduces complication during acquisition, as relatively flat samples can be entirely imaged without the need for autofocus systems or slide tilt/deflection correction.

## 4. Discussion

While many other implementations of multifocal plane microscopy have previously been demonstrated, our optical system is comparatively simple, versatile, and requires no complex or custom optics. As a result, this system is substantially less expensive and easier to construct than alternative multifocal plane microscopy systems. In fact, the total cost of our four-camera detection system (including cameras, optics, and optomechanical components) was approximately a quarter of the price of a typical sCMOS camera. Despite our system’s simplicity and affordability, the image quality of our system is not compromised, as each camera’s image is relayed through a nearly 4-f system (thus preserving telecentricity), and the beam-splitters in the optical system are in infinity space. Due to the recent improvements in CMOS image sensor technology, this simpler optical system was still able to obtain volumetric images of quality comparable or better than previous works, at four times the speed of a single-camera imaging system. We expanded upon previous publications of multi-camera systems by demonstrating a variety of the unique imaging modes possible with a multi-camera system and super-resolution SIM reconstruction algorithms. Though long exposures introduced substantial amounts of hot-pixels in the raw image data, we were able to achieve satisfactory results by implementing a simple hot-pixel removal step in our image processing procedure.

However, this four-camera system does have multiple drawbacks. Primarily, the intensity of light reaching each camera is ¼ of that in a traditional single-camera system. This reduction of light intensity on each detector necessitates longer exposure times (and thus slower imaging) on dim samples, such as the *Drosophila melanogaster* larvae and HEP-G2 cells imaged in this study. Though this downside would be substantial on previous industrial CMOS technology, the high-sensitivity and low-noise image sensors used in this study were able to image such dim samples at reasonable exposure times. Additionally, this light intensity reduction is a downside common to all multifocal plane imaging techniques known to the authors. Another issue present in our setup is the slight brightness variations between the images produced by the four cameras. Such brightness variations cause stripes in orthogonal projections of images acquired using the Z-stack mode, as visible in Fig. 5i. These brightness variations are primarily due to tolerances in the transmission/reflection ratio of the beam-splitters used, which can result in intensity variations of ∼5% between cameras. This issue could be resolved by using alternative beam-splitters with an improved transmission/reflection ratio tolerance. In this study, we were able to minimize these striping artifacts by normalizing the intensity of the image data between cameras after acquisition, though this method is imperfect (as seen in Fig. 5i). Finally, the simple re-imaging system used in our four-camera system introduces slight axial chromatic aberration into the image, an aberration which is difficult to avoid.

Our four-camera system presents significant potential for future work. Firstly, the hot-pixel removal method used in this paper is functional, but primitive. Hot pixels that appear in the data which were not captured in the dark frame are not removed by this simplistic method. More sophisticated hot-pixel removal techniques, such as the one described in [37], could be used instead for higher-quality results.

Furthermore, the four cameras in the system described here were used to image separate focal planes, but the detectors could also be used in slightly different configurations for an even wider variety of imaging types. By aligning the four cameras without defocusing the sensors, the cameras can quickly be triggered in succession to achieve frame rates 4x higher than the cameras are individually capable of. Additionally, if the cameras are focused to the same plane but displaced laterally with the translation stages, the FOV of the detection system can be expanded by up to 4x. This is particularly useful for cameras with small sensor sizes relative to the imaging circle of the microscope, such as the cameras in our setup. Finally, if aligned laterally and kept focused, the four-camera setup could be used to generate high dynamic range (HDR) images by acquiring four images simultaneously with different exposure times, then merging these images using well-known HDR algorithms [38–40]. These potential uses of a multi-camera system have previously been explored in the context of photography [41], but remain mostly novel in the field of microscopy.

Note that all of these additional imaging modes require no adjustment to the optics of our system – all that must be changed is the positioning and triggering synchronization of the cameras. As such, if the translation stages used to position the cameras were motorized, the four-camera detection system could quickly and precisely be reconfigured for a variety of uses. In the context of this work, such a motorized system could allow for quick adjustment of focal plane spacing, as well as to automatically compensate when the microscope objective is changed. The system could also be easily reconfigured into the high frame rate or expanded lateral FOV modes previously discussed without tedious alignment. With sufficient image processing, a motorized system could even automatically align and focus the cameras.

## 5. Conclusion

To summarize, we have presented in this study a highly flexible and practical multiplane imaging system using multiple cameras. This system is effective at quickly obtaining high-quality 3D fluorescence images using the three imaging modes explored in this paper, and we also proposed multiple additional imaging modes that could be explored with the same optical setup. The effectiveness, simplicity, and versatility of our optical system provide a promising approach for future implementations of multifocal plane imaging in fluorescence microscopy.

## Supporting information

Visualization 1

## Acknowledgements

Research reported in this publication was supported by the National Institute of General Medical Sciences of the National Institutes of Health under award No. 1R15GM128166–01. This work was also supported by the UCCS BioFrontiers Center. The funding sources had no involvement in study design; in the collection, analysis, and interpretation of data; in the writing of the report; or in the decision to submit the article for publication.

